# Advancing Great Lakes Coastal Wetland Food Web Models using an Integrative Tracer Approach

**DOI:** 10.64898/2026.05.27.725968

**Authors:** Alyssa M. Smith, Matthew J. Cooper, Ryan Otter

## Abstract

Coastal wetlands of the Laurentian Great Lakes support abundant populations of fish, invertebrates, and vegetation, though the trophic linkages connecting primary production and lower consumers is not well understood in these systems. We implemented a multiple-tracer approach to evaluate trophic pathways, pairing traditional food web isotope tracers like carbon (δ^13^C) and nitrogen (δ^15^N) with total mercury concentrations (THg). We predicted that filamentous algae would be the dominant energy resource in the diet of lower trophic-level invertebrates in the Grand River Estuary, a network of riverine coastal wetlands adjacent to Lake Michigan. In addition, we predicted that adding THg as a tracer would improve the resolution of our food web models by clarifying trophic levels and relationships between wetland species. Four basal energy sources were sampled, including filamentous algae, emergent macrophytes, submersed macrophytes, and phytoplankton, along with organic detritus. Aquatic invertebrates were sampled across multiple functional guilds to represent primary and secondary consumers and included amphipods and odonates. Our findings suggest that organic detritus is the dominant resource responsible for energetically supporting these lower trophic levels in the Grand River estuary, although submersed macrophytes were important alternative energy sources for secondary consumers. THg concentrations enhanced the resolution of dietary contribution estimates in MixSIAR models applied to consumer and source data. Isotope biplots revealed that THg concentrations were a more reliable predictor of trophic position than δ^15^N in Grand River Estuary (GRE) sites. This methodology has important implications for future food web studies in complex ecosystems such as coastal wetlands and demonstrates the novel use of mercury as an ecological tracer in a Bayesian mixing model approach.

## Introduction

Lindeman’s (1942) foundational “trophic-dynamic” viewpoint of community ecology emphasized trophic interactions as a crucial element of understanding species interactions and community dynamics and continues to inform modern approaches to food web modeling and analysis. Food webs are networks of trophic relationships that provide graphical representations of nutrient and organic matter exchange from basal energy sources to consumers (Wilbur 1997; Thompson et al. 2012; Krumins et al. 2013). Understanding trophic interactions and the flow of energy within complex ecological networks remains a fundamental challenge within ecology (Sutherland et al. 2012), as trophic dynamics are central to understanding broader ecosystem processes. In aquatic ecosystems, the large number of trophic relationships and species interactions as well as stressors such as climate change and anthropogenic disturbances result in complex food webs (Garcia-Oliva & Wirtz 2025). In aquatic food webs, primary consumers such as macroinvertebrates and zooplankton are fueled by primary production from aquatic vegetation and algae, and energy and carbon are transferred to higher trophic levels via predation and are frequently recycled during this process. Given the inherent complexity of trophic dynamics within aquatic ecosystems, there is a growing need for integrative tracing approaches to accurately quantify energy flows and trophic pathways.

Stable isotopes, trace metals, and fatty acids can provide valuable insight into the flow of nutrients and energy through a food web (Hebert et al. 2006). Food web tracers are widely used to evaluate trophic relationships and provide dietary information. Such tracers characterize organic tissue and are often a more versatile and time-integrated approach to food web analysis compared to simpler methods such as identifying and quantifying gut contents, which offer only a brief snapshot of feeding behavior (Soto et al. 2013; Eglite et al. 2023).

Stable isotope analysis is one of the most widely used approaches in food web research for resolving trophic interactions and tracing energy flow within a food web (Boecklen et al. 2011; Eglite et al. 2023). Stable isotope analysis has been effective in evaluating diet composition, resource utilization, niche properties, food web construction, and feeding and foraging ecology of organisms within a community (Fry et al. 2006; Middelburg 2014; Boecklen et al. 2022). This methodology is based on the use of stable isotope composition in organic tissues, which represent the ratio of the heavy form of an isotope to the lighter form. With stable isotope tracers such as ^13^C and ^34^S, the stable isotope composition of a consumer’s tissues will reflect the composition of its diet, resulting in a conserved signal that can be traced through a specific trophic pathway (Boeklen et al. 2011). Alternatively, tracers such as ^15^N isotopes can be used to estimate trophic position (Peterson & Fry 1987) due to a predictable enrichment of approximately 3.4‰ per trophic level. This occurs as a result of fractionation, a process by which ^14^N is preferentially excreted, resulting in a greater proportion of ^15^N available to be incorporated into consumer tissues. The combined use of carbon and nitrogen isotopic compositions in organic tissues represents the most common stable isotope analysis approach in food web research (Middelburg 2014).

Although stable isotope analysis is an effective tool in dietary analysis and food web modeling, there are limitations to this methodology that arise from such issues as uncertainty in fractionation rates in some ecosystems and overlapping ^13^C stable isotope values among producer groups (Boecklen et al. 2011; Soto et al. 2013). Alternative approaches that incorporate trace elements as tracers can be effective and reliable methods for identifying key trophic pathways (Soto et al. 2013). Although trace metal bioaccumulation has been thoroughly studied in aquatic food webs in recent years, it has rarely been used as a food web tracer (Stewart et al. 2004, Soto et al. 2013), despite its capacity to resolve limitations of SIA, such as overlapping stable isotope values among basal resources.

Mercury is a bioaccumulative metal that increases in concentration as it is transferred to higher trophic levels via consumer diets (Ratkowsky et al. 1975). Mercury naturally occurs in two forms, including inorganic elemental Hg and organic methylmercury (MeHg). Total mercury (THg) represents the combined organic MeHg and inorganic Hg. Upon methylation, a process driven by microbes that are often abundant in marine and freshwater anaerobic sediments, Hg is converted into its organic form (Parks et al. 2013; Ma et al. 2019). Additionally, the conversion of Hg into MeHg provides the basis for the biomagnifying property of mercury within food webs. The biomagnification of mercury, like other trace metals, occurs among consumers because the dietary uptake of THg is greater than elimination, resulting in increased concentrations relative to their prey (Soto et al. 2013) and enabling its use as a tracer to approximate the trophic position of organisms. Cabana et al. (1994) demonstrated a positive correlation associated with □^15^N and mercury concentrations in aquatic food webs using Lake Trout, highlighting its potential as an informative tracer that may supplement the standard carbon-nitrogen approach. Although historically only studied in a toxicological context (Soto et al. 2013), this bioaccumulative property of mercury can be effective when evaluating trophic pathways in aquatic food webs, particularly those in freshwater wetlands, where a simple stable isotope approach may not provide sufficient resolution of trophic relationships and dietary sources.

Wetland food webs are historically challenging to resolve due to the vast number of potential basal energy sources and various physical factors occurring at multiple temporal and spatial scales that drive biodiversity in wetland habitats (Keough et al. 1996; Keough et al. 1999; van der Merwe & Hellgren 2016). In freshwater wetlands, detritus represents a key resource that is isotopically heterogenous due to its highly variable mixture of decomposing biomass. This presents a particular challenge in food web resolution, since detritus is ubiquitous and may likely be a dominant source of energy to lower trophic levels (Moore et al. 2004, Layer et al. 2013). Traditional approaches such as gut contents analysis are unable to provide a time-integrated perspective into dietary sources (Jones & Waldron 2003; Eglite et al. 2023) and simple bulk stable isotope analysis is often ineffective since sources may overlap in isotope signatures based on similarities in carbon-fixation pathways (Keough et al. 1996).

In the Great Lakes region, freshwater coastal wetlands are shielded from wave energy and longshore currents, enabling the extensive development of both emergent and submersed rooted vegetation that can thrive in these sheltered habitats (Keough et al. 1999). This allows for the accumulation of nutrient-rich, organic sediment in the benthic zone (Albert et al. 2005). Open-water areas with access to direct sunlight allow proliferation of algal metaphyton communities. These abundant communities of vegetation trap nutrients and sediment within the ecosystem; an ecosystem service which provides protection to nearby bodies of water and drives primary productivity (Herdendorf et al. 1987; Jude et al. 2005). These freshwater coastal ecosystems often house diverse and productive algal, macrophyte, and aquatic macroinvertebrate populations that provide energetic resources to consumers such as migratory waterfowl and fish (Cooper & Uzarski 2016). Great Lakes river mouth wetlands in particular host diverse aquatic macroinvertebrate communities that thrive due to the system’s high productivity (King & Brazner 1999, Cooper et al. 2006). Knowledge about food web dynamics in freshwater coastal wetlands remains limited, though past research has established important trophic linkages mediated by mobile fish consumers connecting nearshore and littoral habitats in Great Lake and coastal wetland habitats (O’Reilly et al. 2023). Other studies have demonstrated the mutual energetic subsidy in food webs within coastal wetlands and Great Lake nearshore habitats (Sierszen et al. 2019), as well as the abiotic factors driving biodiversity and isotopic niche plasticity in coastal wetlands (Keough et al. 1999; Sierszen et al. 2012, Rojas et al. 2025). Yet, quantifying the structure of Great Lakes coastal wetland food webs and the major trophic pathways linking primary and secondary production remains a challenge for ecologists in the region due to the unique nutrient dynamics that govern the cycling of C and N isotopes.

The Grand River Estuary (GRE) is a rivermouth ecosystem that occurs at the interface of Lake Michigan and the Grand River, one of its major tributaries (Fig. 1). Rivermouth ecosystems of the Laurentian Great Lakes are often viewed as analogous to marine estuaries due to the direct surface-water connection with the Great Lake, resulting in coupled hydrologic patterns that give these systems the distinction of “freshwater estuaries” (Trebitz 2006). Hydrologic input from the adjacent Great Lakes into estuaries plays a major role in mediating the dynamics of these ecosystems due to its effects on vegetation community structure (Keough et al. 1999; Trebitz et al. 2002, Anderson et al. 2023) and water chemistry (Trebitz et al. 2005). Seasonal changes in water level occur due to snowmelt and greater precipitation in the spring, while fluctuations over longer periods (i.e., years to decades) are caused by broader spatial and temporal patterns in precipitation and evaporation (Gronewold et al. 2013). These hydrological patterns have a profound influence on the ecology of coastal wetlands in the region and drive patterns in biodiversity of vegetation and wildlife (Herdendorf 1990; Anderson et al. 2023). These wetlands are distinct from inland wetlands of the Great Lakes basin as a result of their surface water connection to the nearby lake, which permits their hydrologic and chemical linkages (Brazner & Trebitz 2016). The Grand River watershed, located in southwestern Michigan’s lower peninsula, spans an area of approximately 8972 km.

**Figure 1:**
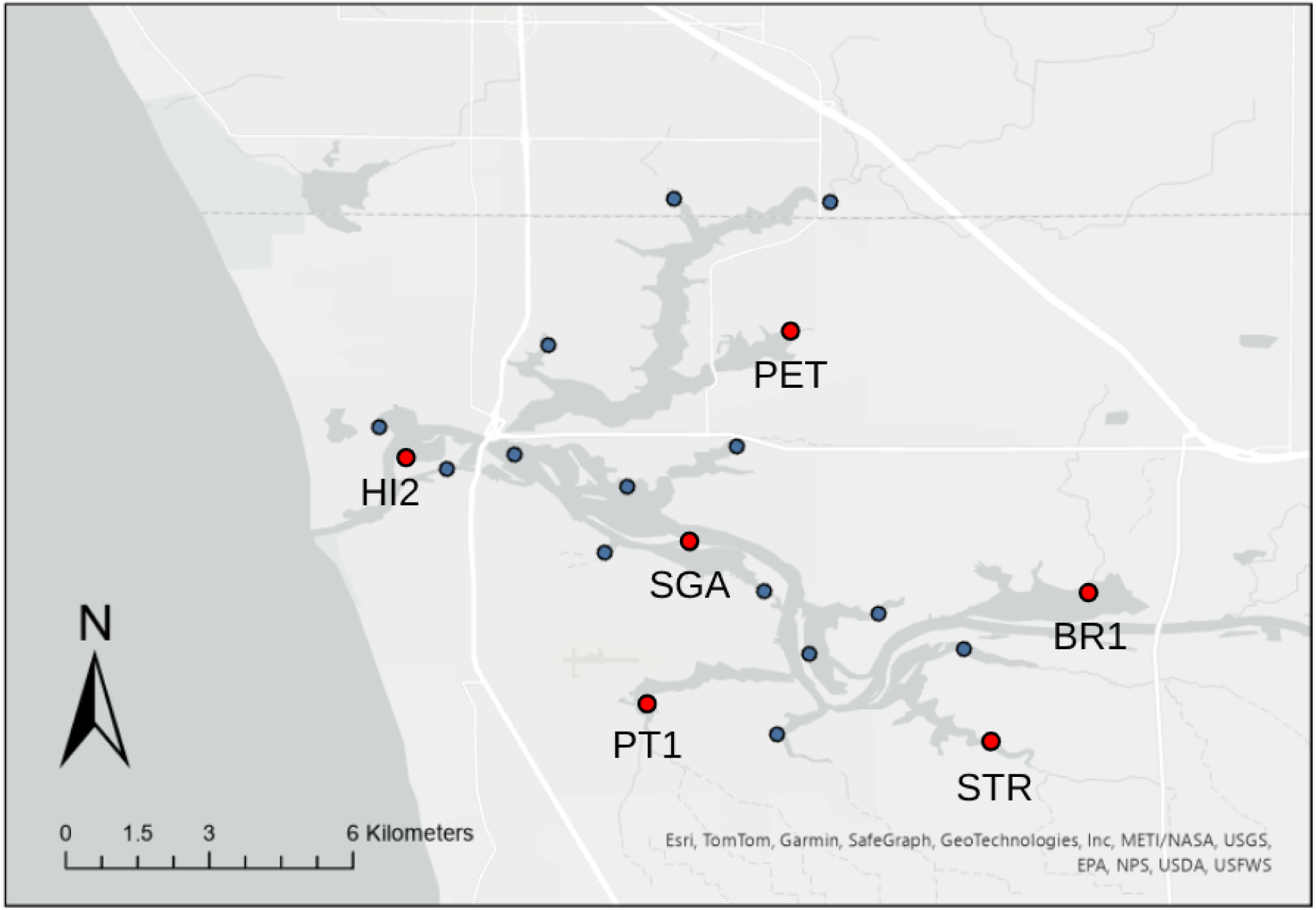
Map of wetland sites in the Grand River Estuary (GRE) with selected sites highlighted with a red point, including Harbor Island (HI2), State Game Area (SGA), Petty’s Bayou (PET), Pottawattomie Bayou (PT1), Stearns Bayou (STR), and Bruce Bayou (BR1). This map was created using ArcGIS® software by Esri.

The aim of this study was to determine whether trace mercury concentrations can be used to identify key trophic pathways and basal energy resources, and develop an approach using total mercury data to modeling lower trophic level food webs in a freshwater estuary ecosystem. Historically, estuarine consumers were thought to be predominantly fueled by macrophyte detrital biomass entering food webs via detritivory and the microbial loop (Odum 1971; Teal 1962), although recent studies suggest that algae may have a more critical role in invertebrate diets due to their digestibility, high rates of production, and preference by consumers (Guo et al. 2016, Hart & Lovvorn 2003; McCutchan & Lewis 2002). We predicted that filamentous algae-derived energy would be a key trophic pathway for lower trophic-level consumers among GRE wetlands, especially invertebrates belonging to grazer and collector-gatherer functional guilds. We included predaceous invertebrates in our study to examine the relative importance of such dietary sources to a higher trophic level. In addition, we predicted that incorporating THg concentrations in food web models would improve their resolution, supplementing the standard C and N stable isotope approach.

## Methods

### Field Sampling

Wetlands within the Grand River Estuary (GRE) form an expansive and connected network (Fig. 1). that can be classified as riverine or drowned river mouth wetlands due to their distinct morphologies (Albert et al. 2005). Six wetlands from the GRE were selected for this study based on site accessibility and habitat characteristics including depth, distance from the river mouth, and presence of key vegetation and invertebrate taxa to ensure that trophic representatives were present at each site. Within each wetland site, three replicate sampling stations were selected within a roughly 300-500 meter transect to conduct all macroinvertebrate and vegetation sampling. Water chemistry and environmental data recorded at each site included pH, dissolved oxygen (DO), total dissolved solids (TDS), specific conductivity (SPC), water temperature, depth, and organic sediment depth. Water chemistry parameters were recorded using a multi-parameter YSI probe (Yellow Springs Instruments Pro Plus Sonde).

At each replicate sampling station, we obtained a sample of submersed aquatic vegetation, emergent macrophytes, and algae (filamentous and planktonic) by collecting material by hand or with nets at the surface. These sources represented the dominant primary producer types in the GRE. Sampled emergent vegetation included water lily (*Nymphaea odorata*) and yellow pond lily *(Nuphar advena*). Submersed vegetation was dominated by Eurasian watermilfoil (*Myriophyllum spicatum*) and hornwort (*Ceratophyllum demersum*), and algae samples were represented by a combination of mixed filamentous algae (primarily *Cladophora* and *Spirogyra*) and duckweed (*Lemnoideae*). Duckweed was included with algal samples because these taxa were often growing as a complex at the water surface and their isotope signals were presumed to be similar because duckweed is not rooted in sediment and floats on the surface of water, similar to metaphyton. Invertebrates were collected using a D-framed dip net to sample from the water column or to remove organisms from the surface of emergent vegetation. Invertebrate samples spanned two presumed trophic levels based on functional feeding guilds, including amphipods (collector-gatherers) and odonates (predators). Amphipods comprised our second trophic level (primary consumers) and odonates represented our third trophic level (secondary consumers). Multiple specimens within each consumer category were collected at each sampling location and combined to increase overall mass for analysis. Collected amphipods included specimens from families including Hyalellidae and Gammaridae, and odonates included specimens from Coenagrionidae, Aeshnidae, and Libellulidae. For most consumer taxa, a range of 2-5 individuals were pooled into composite samples. The number of individuals pooled into a sample varied among taxa and sampling sites because invertebrates exhibited a wide range of size and mass. The sampled invertebrates were generally small, short-lived organisms, which enabled us to use composite samples without compromising the accuracy of isotopic data given that the isotopic compositions of these organisms typically exhibit low variability among individuals and sample pools are advantageous for “pre-averaging” (Fry, 2006). Suspended particulate organic matter (POM) samples were obtained by collecting water samples in triplicate at each site and filtering water through a three-chamber sampler with mesh chosen to retain zooplankton (200 μm), mixed seston (63μm), and phytoplankton (10μm). Benthic organic detritus was collected at each replicate sampling station using a Petit Ponar dredge. Dredged organic detrital material was emptied into a plastic tote and samples were collected from the surface (top 5 cm) of the dredged material. All organic sample types were stored at -18°C prior to laboratory processing.

### Sample Processing

All organic samples except detritus and POM were rinsed with deionized water prior to drying. Vegetation, invertebrates, and detritus were dried in aluminum weigh boats at 60°C for at least 24 h. POM samples were centrifuged at 3000 rpm for ten minutes, excess supernatant was removed, then samples were dried at 60°C for at least 24 h. After drying, all samples were ground into a fine, homogeneous powder using a mortar and pestle.

### Data Collection

Isotopic analyses for δ^13^C and δ^15^N were conducted at the University of Arkansas Stable Isotope Laboratory (UASIL) in Fayetteville, Arkansas using a CN analyzer (EA Isolink, Thermo Fisher) and isotope ratio mass spectrometer (Delta V Plus, Thermo Fisher). All isotope values are reported as ratios of heavy and light isotopes relative to international standards using known reference values: Peedee Belemnite is used for C, and atmospheric nitrogen is used for N. Ratios for each isotope are expressed as parts per thousand relative to these standards using the following formula:

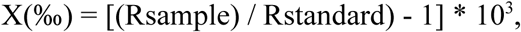

where X is ^13^C or ^15^N, and R represents the ratios of C (^13^C/^12^C) or N (^15^N/^14^N).

Samples were analyzed for THg using US EPA method 7473 on an AGS Scientific MA-3000 direct mercury analyzer located at the Annis Water Resources Institute (AWRI) in Muskegon, Michigan. We could not obtain THg measurements for phytoplankton due to insufficient sample mass.

### Statistical Analysis

Habitat data were evaluated for site-specific differences using an analysis of variance (ANOVA). All habitat data were distributed normally based on a Shapiro-Wilk test with the exception of SPC, which was log-transformed prior to analysis. A Spearman correlation was used to evaluate the relationship between THg and ^15^N because data did not meet the assumption of normality. Kruskal-Wallis and Mann-Whitney U tests were used to compare THg and ^15^N across three trophic levels because isotope data violated parametric assumptions and contained outliers. A trophic position variable was added to this dataset, with trophic level (TL) 1 indicating primary producers, TL 2 indicating primary consumers (amphipods), and TL 3 indicating secondary consumers (odonates) based on known functional feeding guilds (Merritt, Cummins, & Berg 2017; Thorp, Rogers, & Covich 2015). This variable was used to explore the relationship between THg concentrations, δ^15^N, and trophic level to determine whether these values increased with trophic position.

All isotope biplots were generated in R (version 4.4.2) using the car and ggplot2 packages (Fox & Weisberg 2019; Wickham 2016). Bayesian isotope mixing models were fitted using R package MixSIAR (Stock & Semmens 2016; Stock et al. 2018) in R version 4.4.2 (R Development Core Team 2023). This modeling framework integrates a set of parameterizations that improve upon the error structure of previous iterations of Bayesian mixing models, such as MixSIR and SIAR (Stock et al. 2018). We selected this framework due to its capability to account for non-isotopic tracers such as THg. MixSIAR models assess possible combinations of sources and identify the most probable combinations that best reflect consumer tracer signals. MixSIAR models run by using three parallel Markov chain Monte Carlo (MCMC) chains with a length of at least 100,000 iterations that simulate the posterior distribution. The results of the applied models are reported as medians and 5-95% credible intervals of dietary proportions from each source. When determining relative contributions of each source, MixSIAR creates a Just Another Gibbs Sampler (JAGS) file to fit the model (Stock & Semmens 2016). Models are evaluated by two diagnostic tests. The Gelman-Rubin test assesses the convergence of the MCMC chains by comparing the variance among the parallel chains. These should be near 1 at convergence, and all values should generally be below 1.1 (Gelman et al. 2014). The Geweke test (Geweke, 1992) evaluates the burn-in period, or an early proportion of the MCMC chain that does not converge to the target distribution. It compares the mean of this burn-in period with the mean of the second part of the chain, which should be similar at convergence (Stock & Semmens 2016). The model estimates probability density functions of variables of interest, for example, the proportion of amphipod diets that include macrophytes. The Geweke diagnostic specifies that the number of variables found to be ±1.96 at convergence is expected to be under 5%. The number of variables within each chain that violate this assumption are reported in the diagnostic. Models are also evaluated based on the convergence metrics provided in the MixSIAR outputs.

MixSIAR requires three major inputs as separate data files; mixture data that describes the tracer profile of consumers, source data to describe tracer profiles of all possible sources used in the model, and discrimination data that contains the trophic discrimination factors for each of the tracers used. Two iterations of these models were produced. In the first, we examined dietary source proportions to consumers without mercury, given that mercury data were not available for phytoplankton. Primary producers that were assessed in the first iteration of models included detritus, phytoplankton, and submersed vegetation. Filamentous algae and emergent macrophytes were excluded due to their low relative importance in preliminary models, and due to the limitations of the number of sources resulting from using only two tracers. In the second iteration, mercury was incorporated as an additional tracer. Producers in these models included filamentous algae, detritus, emergent macrophytes, and submersed vegetation. Phytoplankton were excluded because we lacked mercury data for phytoplankton. Bayesian mixing models are capable of identifying trophic pathways or energy derived from a source to a consumer that may not have directly consumed it. Since a signal from □^13^C and other dietary tracers will be conserved as it passes through multiple trophic levels, a trophic pathway from a basal energy source to a higher consumer can be identified, even if tracer data are not available for other consumers in the same pathway since a source signal will still be represented in higher consumer tissues (Peterson & Fry 1987). Food web studies using Bayesian mixing models typically aim to determine contributions to the diet of a single population of animals. In this case, we aimed to estimate the energetic contributions of sources on a broader scale, thereby requiring a comparison of the three versions of this model, highlighting the relative source contributions to each consumer group. Trophic Discrimination Factors (TDFs) of 1‰ and 3.4‰ were used for □^13^C and □^15^N, respectively (Vander Zanden & Rasmussen 2001). Because the THg data in this study were raw concentrations rather than isotopic ratios, estimating a mathematically ideal TDF for our sources posed a challenge. While stable isotope discrimination factors are derived from predictable physiochemical processes such as fractionation, no universal discrimination factor exists for raw THg concentrations. Therefore, we treat this THg discrimination factor as a calibration parameter to enable the tracer to be geometrically compatible within isospace, rather than a discrete biological trophic discrimination factor. Different factors at varying orders of magnitude (1, 10, 100) were tested through a series of JAGS model runs to evaluate model sensitivity to changes in discrimination factors which were compared based on their DIC, Geweke statistic, and Gelman-Rubin statistic. A trophic discrimination factor of 100 was implemented for THg in our mixing models. Models run using this factor generated the most favorable model diagnostics and demonstrated a good model fit when compared with other factors.

With MixSIAR, the number of sources may not exceed the number of tracers + 1 without resulting in errors during source contribution estimation. This enabled us to use four dietary sources (algae, emergent macrophytes, submersed macrophytes, and detritus) with three tracers (δ^13^C, δ^15^N, THg) in our modeling framework. Though three size classes of POM including zooplankton, mixed seston, and phytoplankton were sampled in the study, these data are only included in biplots and not JAGS models due to limitations in the number of tracers used. Past research has supported that both filamentous algae and macrophyte-derived detritus are major energy sources for aquatic macroinvertebrates (Hart & Lovvorn 2003; Odum 1971). Therefore, we prioritized the four selected producers in our food web models to highlight those specific pathways.

## Results

### Habitat Characteristics

Physical parameters measured at each wetland site include water temperature, depth, pH, dissolved oxygen (DO), total dissolved solids (TDS), and specific conductivity (SPC) (Table 1). The mean water temperature across sites was 23.15°C ± 1.97. Mean %DO across sites was 50.42 ± 29.69 while mean DO (mg/L) was 5.70 ± 1.86. Mean SPC across sites (μS/cm) was 649.05 ± 466.40 while mean TDS (mg/L) was 0.30 ± 0.07. Mean pH across sites was 7.74 ± 0.25 and mean depth (cm) was 58.88 ± 19.45.

**Table 1:**
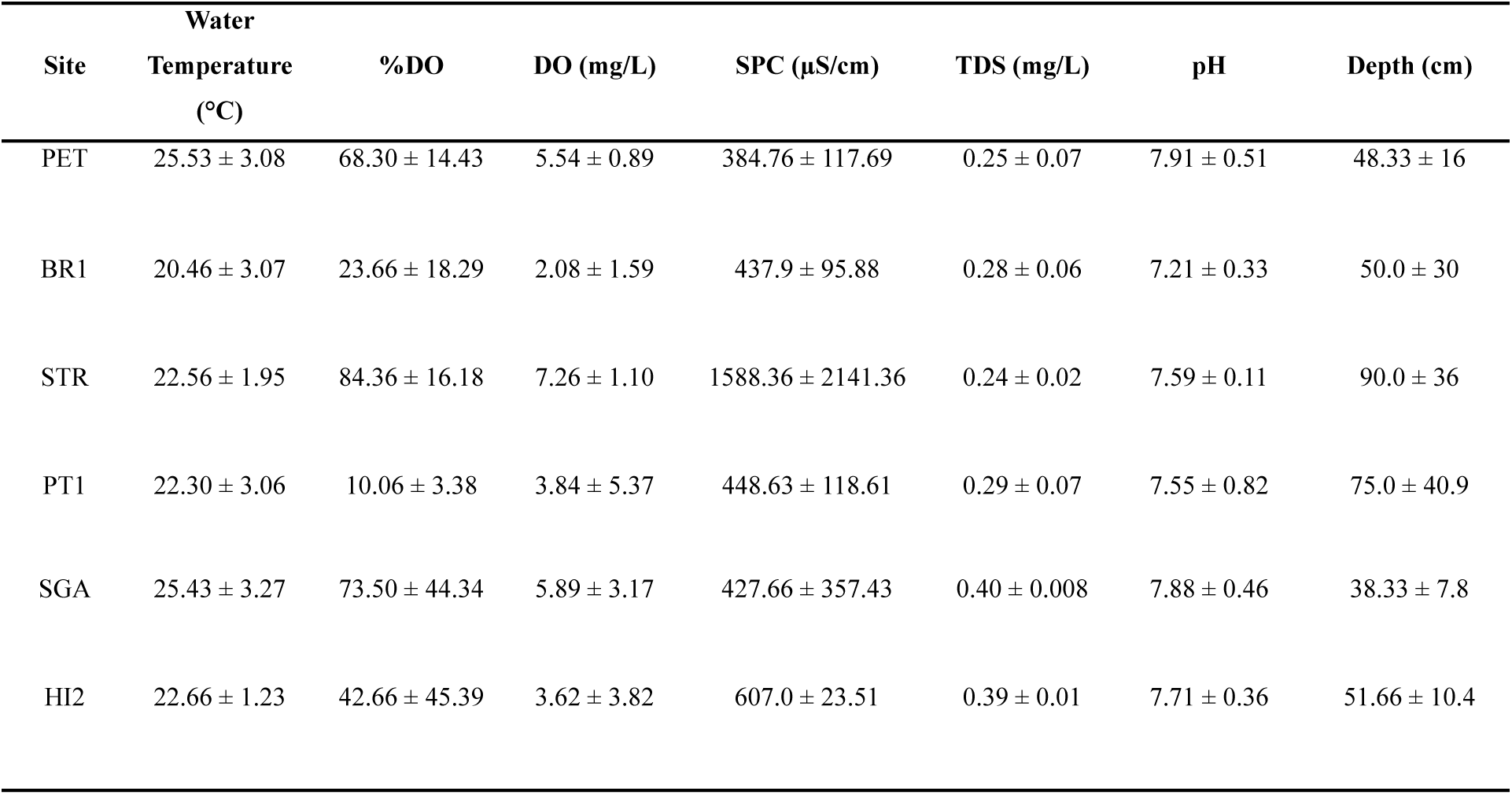
Physical and water chemistry data (mean±SE) across six wetland sites, including Petty’s Bayou (PET), Bruce Bayou (BR1), Stearns Bayou (STR), Pottawatomie Bayou (PT1), State Game Area (SGA), and Harbor Island (HI2).

### Isotopic Compositions

Emergent macrophytes were consistently enriched in δ^13^C relative to submersed macrophytes (Fig. 2, Table S1). Detritus was generally enriched relative to algae, except at BR1 and SGA, and was depleted relative to emergent macrophytes except at SGA and HI2; it was however enriched relative to submersed macrophytes at all sites except BR1. Zooplankton was typically depleted relative to mixed seston and phytoplankton, with PET and BR1 as exceptions, where it was enriched relative to mixed seston and enriched relative to phytoplankton at PET. Mixed seston was depleted relative to phytoplankton at all but PT1. Phytoplankton was enriched relative to zooplankton and mixed seston at BR1, STR, PT1, and SGA. Amphipods were consistently enriched relative to odonates. Across producers and consumers, δ¹³C values at SGA and HI2 were elevated compared to upstream sites.

**Figure 2:**
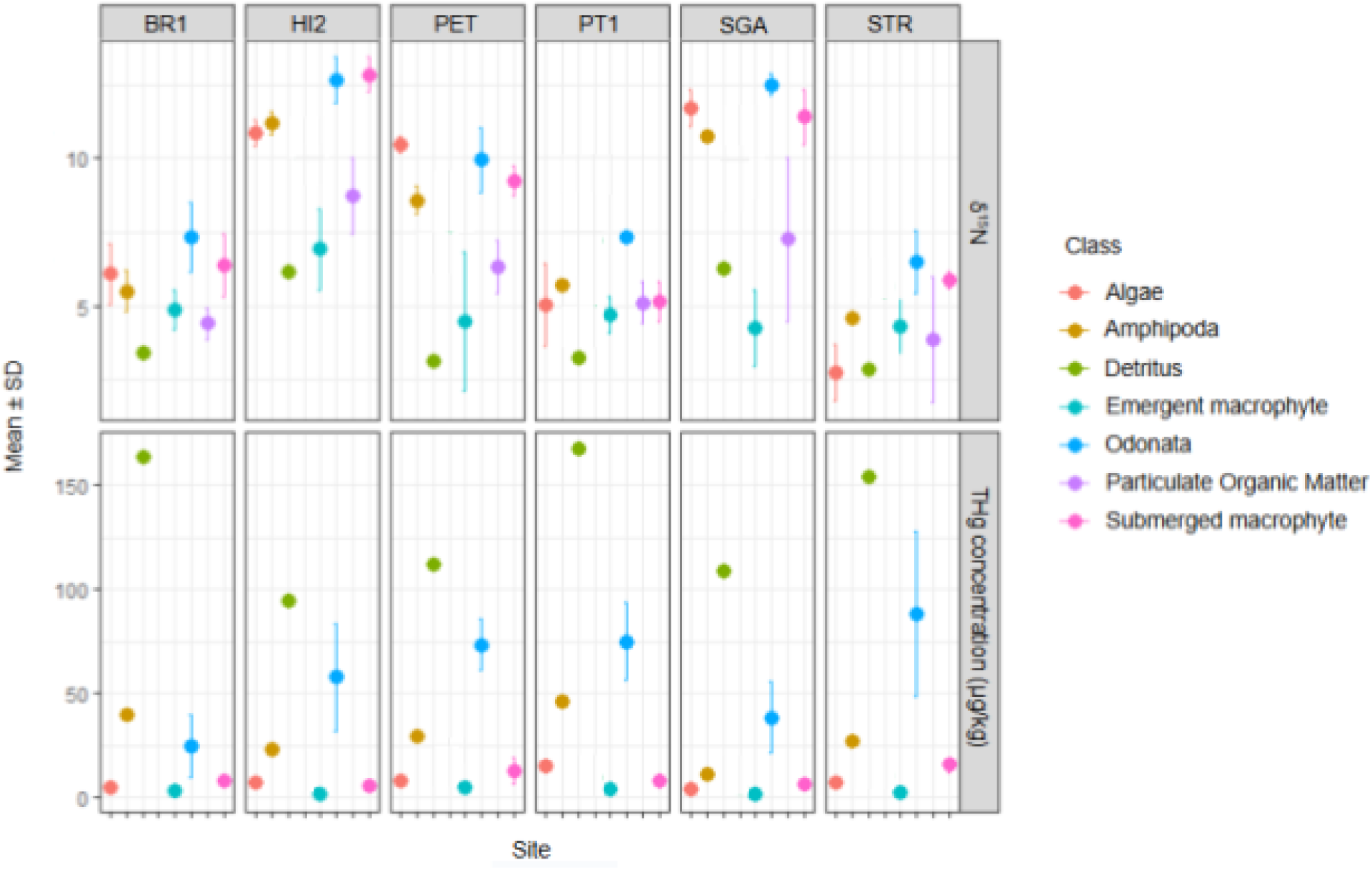
Mean ± SD plots describing δ^15^N and THg concentrations across six wetland sites.

Vegetation exhibited a wide range of variation in ^15^N isotopic ratios across sites (Fig. 2, Table S1). Algae exhibited the most variation at STR, PT1, and BR1. Emergent macrophyte ^15^N ratios varied the most at HI2, PET, and SGA, while submersed macrophytes expressed the widest variability at BR1, PT1, and SGA. The trophic position of algae and submersed macrophytes based on ^15^N ratios varied extensively, presenting a challenge in trophic position approximation using only δ^15^N. Algal δ^15^N was depleted relative to submersed and emergent macrophytes at STR, enriched relative to macrophytes at PET and SGA, and enriched relative to macrophytes but depleted relative to submersed macrophytes at HI2. Detrital δ^15^N was depleted relative to algae and submersed vegetation at all sites, and depleted relative to emergent macrophytes at all sites except SGA and HI2. Zooplankton δ^15^N was only enriched relative to mixed seston and phytoplankton at three sites (PET, SGA, HI2), and depleted relative to mixed seston and phytoplankton at one site (STR). Mixed seston was depleted relative to both groups at three sites (BR1, PT1, SGA). Phytoplankton was enriched relative to both groups at three sites (BR1, STR, PT1) and depleted relative to both groups at two sites (PET, HI2). Odonate δ^15^N was enriched relative to amphipods across all sites.

Producer and consumer δ^15^N was enriched at two sites relative to other sites (Fig. 2, Table S1). At SGA and HI2, algae was enriched relative to other sites by 5.5‰ and 6.3‰, respectively. Submersed macrophytes were enriched relative to other sites by 4.7‰ and 6.2‰, respectively. Emergent macrophytes were not enriched at SGA relative to the mean δ^15^N of other sites, however, emergent macrophytes at HI2 were enriched by 2.3‰. Amphipods were enriched at SGA and HI2 relative to other sites by 4.6‰ and 5.1‰. Odonates were enriched at SGA and HI2 by 4.6‰ and 4.8‰, respectively.

In models where THg data were excluded, sources included detritus, phytoplankton, and submersed vegetation. Isospace plots (Fig. 3) captured substantial source overlap between detritus and phytoplankton across the ecosystem, and some overlap was observed between submersed vegetation and phytoplankton. Submersed vegetation was enriched in ^15^N relative to detritus, with phytoplankton data demonstrating ^15^N overlap across both groups. In models where THg data were included, sources included filamentous algae, detritus, emergent macrophytes, and submersed macrophytes. Isospace plots (Fig. 4) demonstrated some overlap between algae and submersed vegetation. Emergent macrophyte ellipses were broad due to variance in ^13^C ratios, and most detritus data fell within macrophyte isospace. Algae and submersed vegetation were enriched in ^15^N relative to detritus and emergent macrophytes.

**Figure 3:**
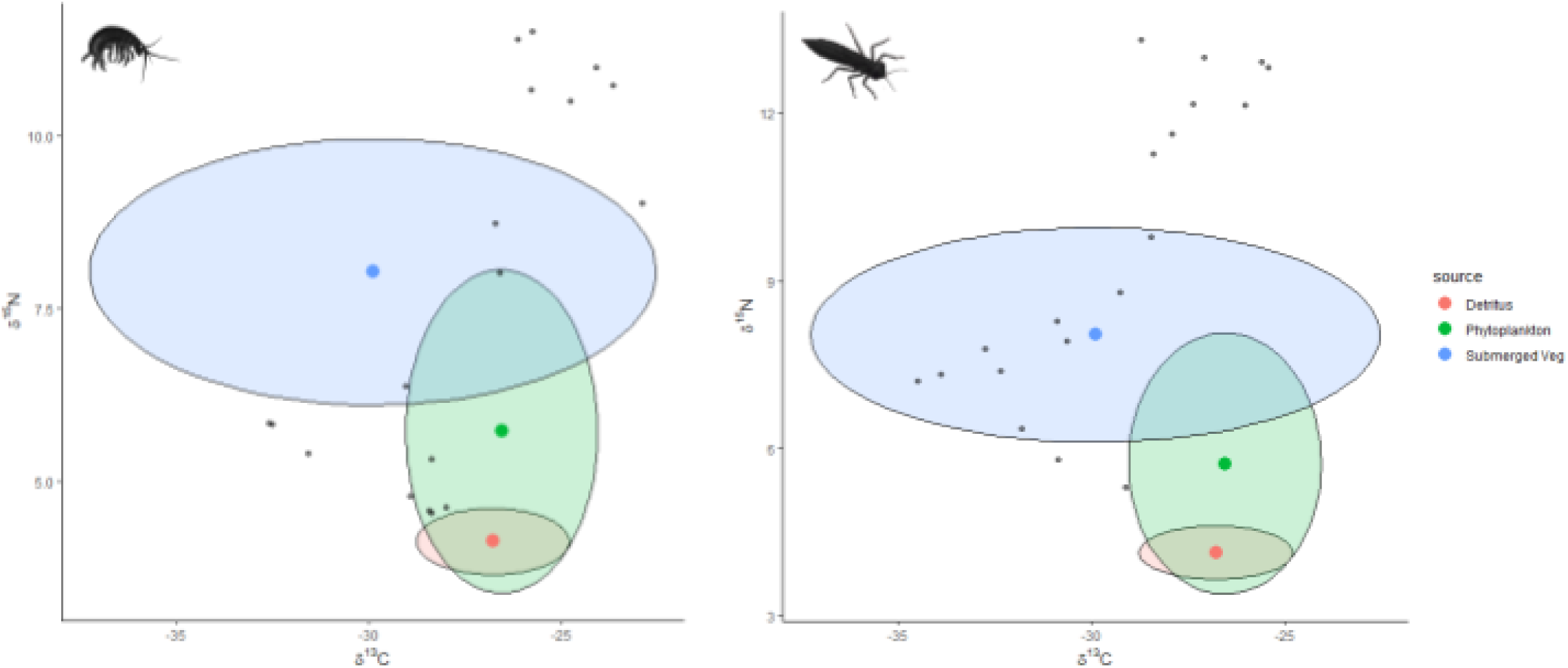
Isospace plots illustrating MixSIAR models comparing source ^13^C and ^15^N data between two consumers. Consumer data is plotted using grey points comparing amphipods (left) and odonates (right). Three sources (detritus, phytoplankton, and submersed vegetation) are included in the models.

**Figure 4:**
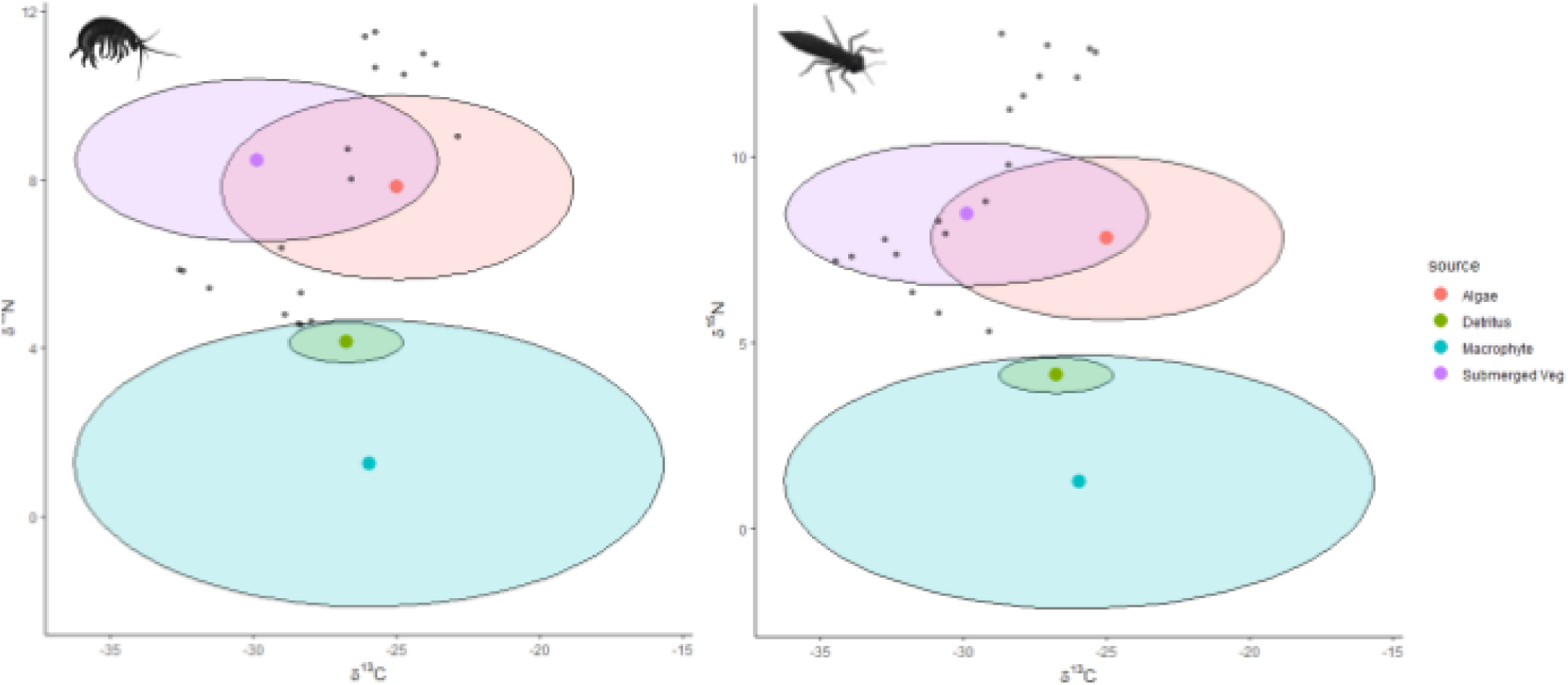
Isospace plots illustrating MixSIAR models comparing source ^13^C and ^15^N data between two consumers. Consumer data is plotted using grey points comparing amphipods (left) and odonates (right). Four sources (filamentous algae, detritus, emergent macrophytes, and submersed vegetation) are included in the models.

### THg Concentrations

THg concentrations of filamentous algae were greater than emergent macrophytes and lower than submersed macrophytes at four sites (Fig. 2; Table S1) (PET, BR1, STR, SGA). Algal THg was greater than both macrophyte types at two sites (PT1, HI2). Emergent macrophyte THg concentrations were lower than other vegetation types at all sites. Detritus THg was greater than all sample types across all sites. An isospace plot was generated to visualize source and consumer data using THg and ^13^C. Detritus exhibited significantly higher THg concentrations than other sources included in the model (Fig. 5).

**Figure 5:**
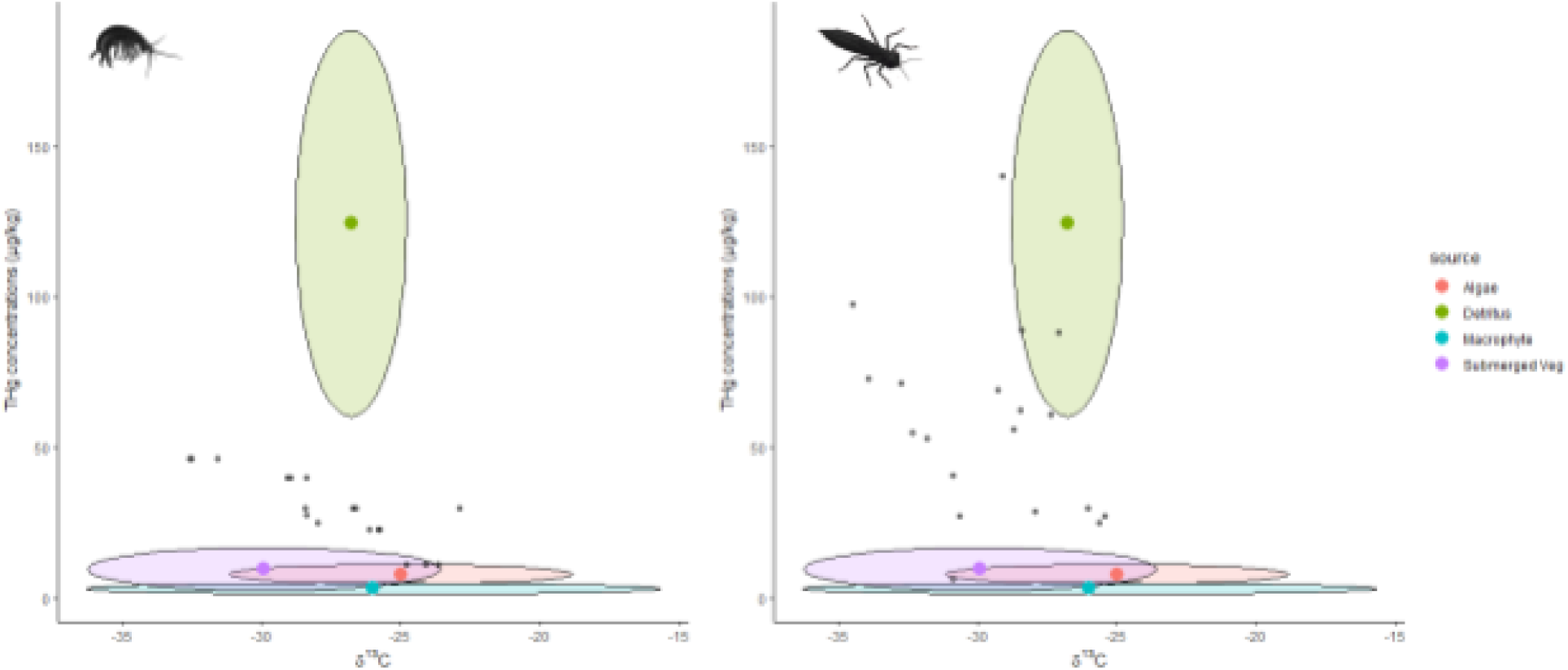
Isospace plots illustrating MixSIAR models comparing source ^13^C and THg data between two consumers. Consumer data is plotted using grey points comparing amphipods (left) and odonates (right). Four sources (filamentous algae, detritus, emergent macrophytes, and submersed vegetation) are included in the models.

### THg and ^15^N

THg concentrations and δ^15^N demonstrated a weak correlation (p = 0.049; r_s_ = 0.21). THg concentrations were significantly different among at least two trophic levels (p = 0.0004) and pairwise comparisons were significant between trophic levels 2 and 3 (p = 0.0001) based on a Mann-Whitney U Test. Comparing ^15^N among trophic levels also revealed a significant difference (p = 0.001), with pairwise comparisons demonstrating significant differences between trophic levels 1 and 3 (p = 0.0014) based on a Kruskal-Wallis test. We additionally calculated a THg trophic magnification factor of 2.94 among our producer groups and invertebrates, using our trophic level variable and log-transformed THg concentrations.

### Mixing Models using *^13^C & ^15^N*

Mixing models applied to isotopic and trace Hg data were run with separate models fit to each consumer group. MixSIAR estimated that diets of amphipods were dominated primarily by detritus (Fig. 6), with contributions ranging from 64.5% to 87.2% across all sites. Submersed macrophytes make up a secondary resource across sites, with phytoplankton contributing minimally to amphipod diets. Submersed macrophytes were the main basal energy resource for odonates (Fig. 6), with proportions ranging from 41.9% to 49.5% across all sites. However, detritus was an important secondary energy source to odonates, with contributions ranging from 35.8% to 42%. Phytoplankton represented the smallest proportion of dietary contributions among odonates, however the contribution of phytoplankton-derived energy was greater in odonates than in amphipods.

**Figure 6:**
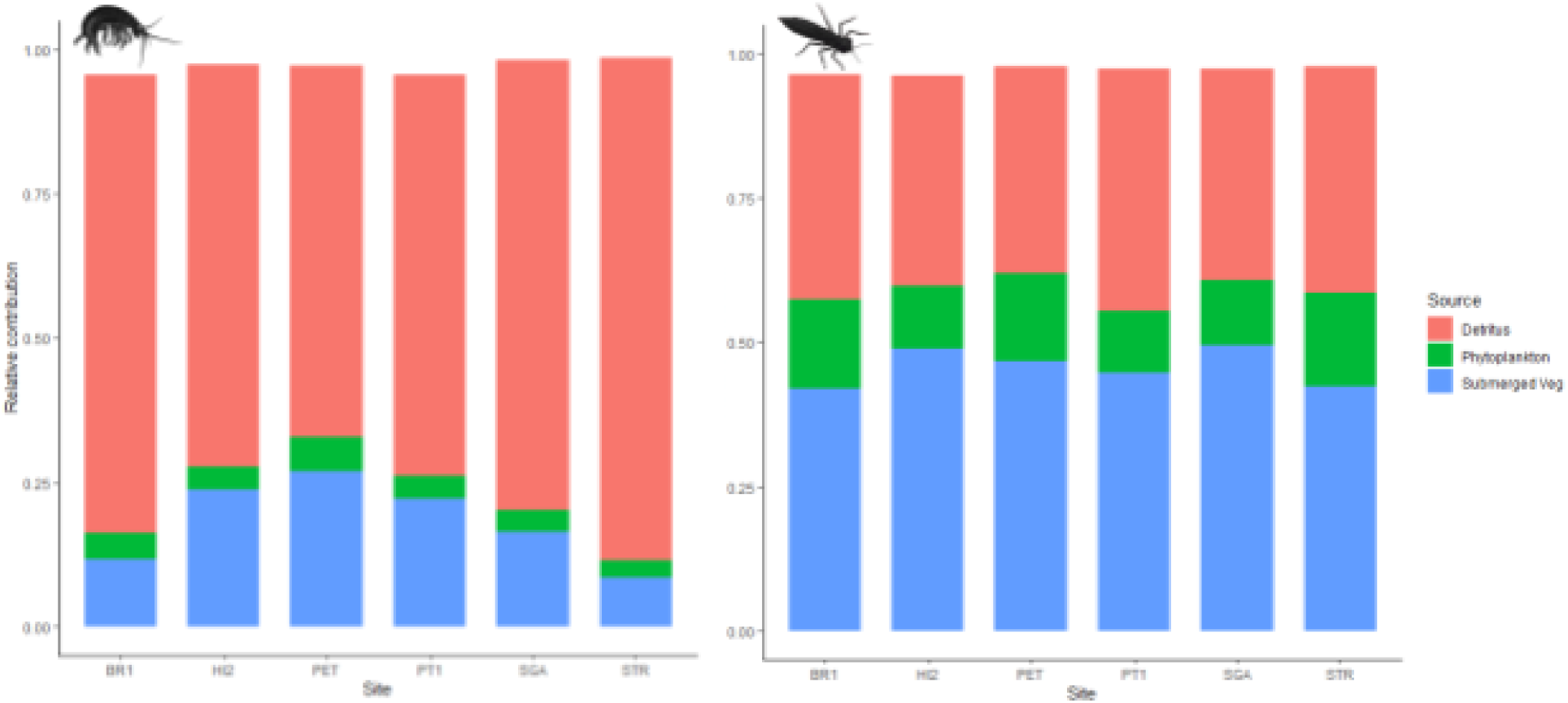
Relative abundance plots demonstrating dietary source contributions to two consumers, amphipods (left) and odonates (right). Sources include detritus, phytoplankton, and submersed vegetation. Plots were generated using source and consumer ^13^C and ^15^N ratios.

### Mixing Models using *THg, ^13^C, & ^15^N*

When THg was incorporated into mixing models as an additional tracer, four sources were evaluated for their contributions to consumer diets. Among amphipods, diets were primarily made up of detritus, with contributions ranging from 40.9% to 59.8% across sites (Fig. 7). Filamentous algae, emergent macrophytes, and submersed macrophytes made important secondary contributions across sites. Submersed macrophytes dominated odonate diets across every site (Fig. 7), with contributions ranging from 54.7% to 63.3% across sites. Both detritus and emergent macrophytes were alternative sources for odonates, however filamentous algae made minimal contributions.

**Figure 7:**
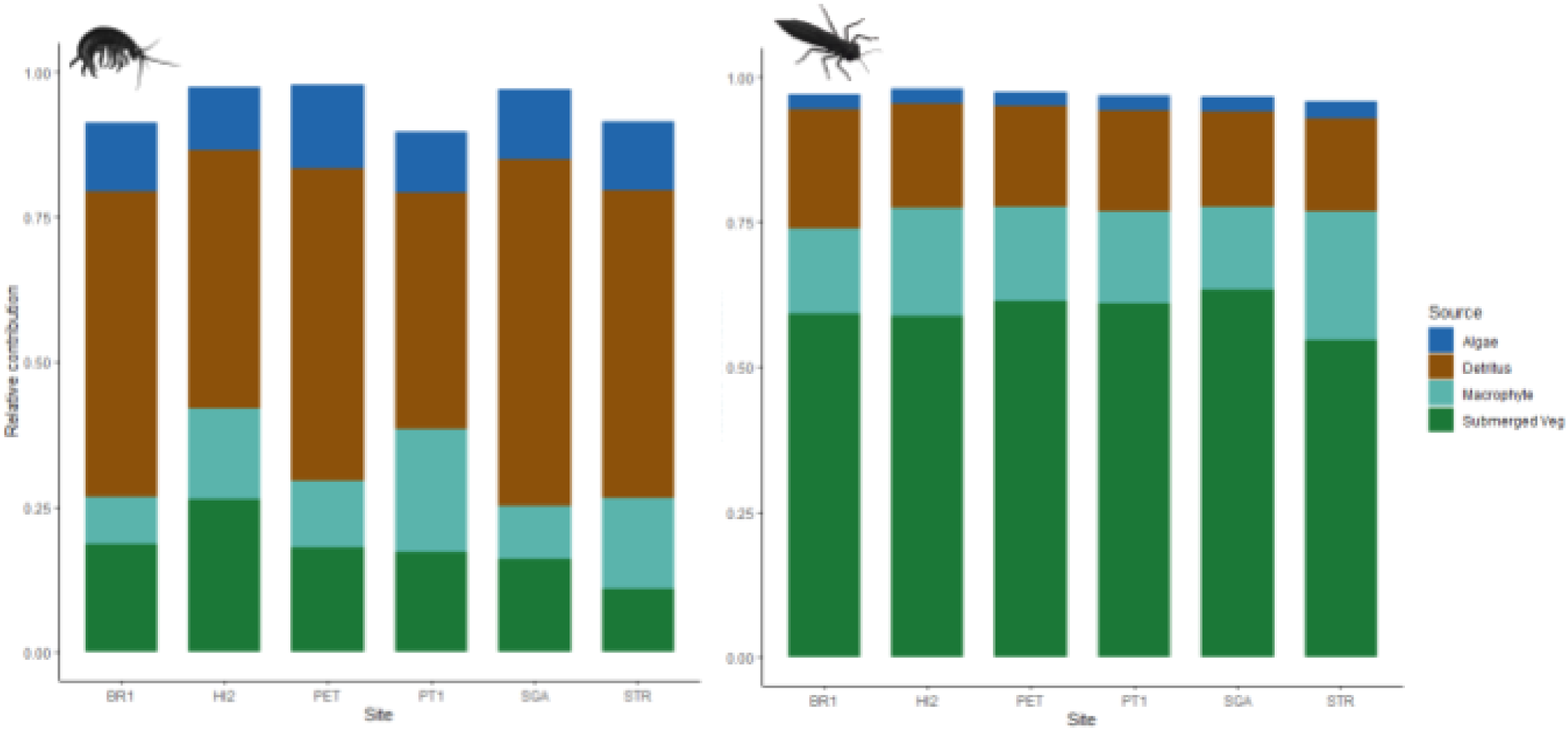
Relative abundance plots demonstrating dietary source contributions to two consumers, amphipods (left) and odonates (right). Sources include filamentous algae, detritus, emergent macrophytes, and submersed macrophytes. Plots were generated using source and consumer ^13^C and ^15^N ratios and THg concentrations.

## Discussion

### Isotopic Compositions

Previous studies have demonstrated that algae exhibits an enriched δ^13^C relative to rooted aquatic plants, due to the differential carbon sources found in their environment (Post 2002). While rooted macrophytes obtain their dissolved inorganic carbon (DIC) from the water column and sometimes from sedimentary bicarbonates, algae, particularly metaphyton, will uptake carbon from both atmospheric carbon dioxide (CO_2_) and other DIC forms from surface water. Assimilation DIC results in isotopically lighter δ^13^C when compared to other forms (Marty & Planas 2008), and the δ^13^C signature of algae and macrophytes reflect the mixture of DIC forms that they assimilate. With the exception of some observed algal δ^13^C depletion relative to emergent or submersed macrophytes at three sites, our data largely supports previous trends outlined in the literature. However, given that some algae had a depleted ^13^C signature relative to macrophytes rather than the expected enriched ^13^C signature, we suspect that algae in some wetlands in the GRE are obtaining a large part of their carbon from an alternate source of DIC such as bicarbonates rather than CO_2_, which may be more bioavailable given the environmental conditions of these wetlands, particularly circumneutral pH (Table 1). Due to the variability observed among producer δ^13^C depletion and enrichment across sites, it is difficult to ascertain whether one source of DIC is preferentially used within each group. Instead, our results suggest high variability in DIC sources in GRE wetlands presumably caused by the rapid cycling of carbon as well as shallow depths enabling increased atmospheric diffusion. This reinforces that a simple C and N tracer approach may not be sufficient when such intense variability exists in the cycling of both elements.

Enriched δ^13^C was generally observed among producers and consumers in the two wetlands nearest to the main channel and the mouth of the Grand River, SGA and HI2. Generally, systems more impacted by urbanization or other anthropogenic stressors will exhibit depletion of ^13^C at the base of the food web rather than enrichment, contrary to what we identify at these sites. While the δ^13^C of producers is largely determined by the δ^13^C of DIC sources, several anthropogenic processes can drive the depletion of δ^13^C in basal resources. A study conducted in tropical systems found an association between the δ^13^C of producers and consumers and the degree of urbanization within the sampled watershed, indicating that more urban land-use was related to depleted δ^13^C among aquatic species (Olsen et al. 2010). The cause of this relationship is undetermined, but may be linked to an increase in terrestrial inputs in these areas (Olsen et al. 2010). Past work also supports that combustion-derived carbon is isotopically depleted in ^13^C and has the capacity to drive the δ^13^C of terrestrial plants sequestering fossil fuel CO_2_ towards depletion (Lichtfouse et al. 2003). Although SGA and HI2 are in close proximity to industrial land use areas, δ^13^C at the base of these food webs does not reflect trends demonstrated in the literature.

Macrophytes and algae exhibited the greatest variation in δ^15^N. This high variability may indicate different sources of N, with either nitrate or ammonium being the most abundant forms. Nitrogen assimilation from different sources can drive fractionation in plant tissues, which may result in the observed variation in macrophytes (Fry et al. 2000). Different metabolic pathways used for N assimilation, availability of N sources, and bacterial denitrification may also be responsible for variation in ^15^N isotopic signatures (Garces-Cuartas et al. 2021; Handley & Raven 1992). Increased nutrient loading from agricultural lands and wastewater effluent is also a possible driver for enriched δ^15^N in both producers and consumers in estuarine systems such as the GRE (Fry 2002). Detrital δ^15^N compositions were depleted relative to other producers, suggesting that detritus may not be entirely composed of macrophyte and algal biomass. While it is likely that organic detritus contains some of this material, we suspect that there are additional inputs to the detrital pool including terrestrial plant biomass. Terrestrial plant δ^15^N is often depleted relative to aquatic plants due to the variety of N sources available to aquatic plants that can result in enrichment (Gong et al. 2021).

Similar to δ^13^C, mean δ^15^N values from both producers and consumers were enriched at SGA and HI2, the two sites closest to the main channel and the mouth of the river. In the early 1950s and 1960s, the area surrounding HI2 (Harbor Island) was used for waste disposal, and later housed a coal-fired power plant where coal ash was disposed of in nearby wetlands (MPART, 2023). SGA (State Game Area) is located about 5 km upstream of HI2. The Grand Haven Wastewater Treatment Plant is located between these two sites. Land use shifts towards urbanization and human activity can alter the δ^15^N of producers and consumers, and on average, δ^15^N is 1‰ greater in more urbanized areas (Steffy & Kilham 2004; Olsen et al. 2010). Urbanization can increase the release of wastewater, resulting in increased rates of nutrient transport to groundwater, making groundwater a key mechanism for delivering wastewater N to estuaries (Valiela et al. 1992; McClelland et al. 1997). This also indicates that ^15^N can act as an indicator of anthropogenic inputs into an estuarine system (McClelland et al. 1997). Given that on average, N was enriched in both producers and consumers at SGA and HI2, we suspect that anthropogenic influence such as agricultural runoff, wastewater effluent or septic inputs to groundwater may be mechanisms driving δ^15^N enrichment in the estuary at these more downstream locations.

### THg Tracer

The use of trophic discrimination factors (TDFs) in recent Bayesian isotope mixing models is crucial to robust food web analysis. This is especially true when used to estimate contributions of basal energy sources to consumers, where appropriate TDFs are necessary to correct isotope values based on trophic fractionation (Phillips et al. 2014). The use of inappropriate TDFs can be a significant cause of error in Bayesian mixing models (Phillips et al. 2014), and are often referred to as the weakest link in mixing model approaches due to model sensitivity (Bond & Diamond 2011). TDFs derived from the literature that apply to some species are not necessarily appropriate for species where the exact factor is unknown (Phillips et al. 2014). Depending on the set of TDFs used in a mixing model, dietary contribution estimates may differ substantially (Diamond & Bond 2011), reinforcing the importance of using appropriate TDFs in diet reconstruction and food web modeling.

Trace Hg concentrations were central to our food web tracer approach in this study, although few data are available for TDFs of THg that are widely applicable to multiple groups of organisms. Hg biomagnification, and therefore a reliable TDF, can vary based on species, environment, and a range of other factors (Lavoie et al. 2013; Sinclair et al. 2024). Accordingly, we selected a TDF that produced the most reliable results that were corroborated by species feeding ecology and provided the best fit model. These tests indicated that a discrimination factor of 100 was best-suited for our mixing models to interpret dietary sources and trophic position within our dataset. In addition, these models describe trophic pathways that are supported by our ecological knowledge of these systems. While this factor provided a good model fit and results supported by known feeding ecology of our consumers, it is likely still imprecise, and further research is needed to address discrepancies in TDFs across different consumer types, ecosystems, and tracers. High variability of δ^15^N, particularly in producer groups, made interpretation of trophic pathways and positions challenging. While mixing models indicated algal, macrophyte, and detrital-derived energy, δ^15^N signals did not reflect the same pathways. Producer δ^15^N signals were enriched relative to consumers, although this did not align with the mixing model results or our assumptions about the relative trophic levels of consumers and producers. When THg concentrations were compared with δ^13^C in biplots rather than δ^15^N (Fig. 2), trophic positions of producers and consumers changed significantly and more closely reflected our assumptions of food web structure. We observed minimal variance in THg concentrations among all sample types (except odonates) but high variability in δ^15^N, suggesting that the use of δ^15^N to estimate trophic position in systems with high ^15^N variability may be unreliable. In similar systems, THg concentrations may be a useful alternative to ^15^N for trophic position approximation, especially for applications such as mixing models.

## Mixing Model

Results from our mixing models revealed several key trophic pathways, including detritus and algae-based pathways for both primary consumers, and detritus and submersed vegetation-based pathways for secondary consumers. Detrital-derived energy has long been thought to be the primary energy source for lower trophic levels in aquatic food webs such as those in the GRE (Odum 1971; Teal 1962, Jeffres et al. 2020). However, representation of algae in aquatic macroinvertebrate diets is strongly supported by the literature (Hart & Lovvorn 2003; McCutchan & Lewis 2002) due to high rates of primary production and consumer preference. Our findings suggest that, while still contributing significantly to consumer diets, algae is a secondary dietary resource relative to detritus for many primary consumers in GRE wetlands. While emergent and submersed vegetation can be an energy resource for invertebrates in these systems, they are also utilized as refuge from predators and as substrate for algal growth (Campeau et al. 1994, Nieoczym et al. 2023, Baattrup-Pederson et al. 2025). Emergent macrophytes were not strongly represented in consumer diets, however, submersed vegetation provided a significant source of energy to odonates. This pathway likely occurs via indirect consumption, given that odonates are predators, and there is likely an intermediary consumer not sampled in our study that provided this energy to odonates. Additionally, it is likely that pre-detrital aquatic macrophyte “litter” may be an important resource for many invertebrates in this study. Lower consumers may be capable of foraging small particles of macrophyte material that settles into benthic sediment in GRE wetlands. Particles of plant matter that have not yet fully decayed and entered the mixture of fine organic detritus will likely retain an isotope signature that is comparable to living macrophyte tissue. This may explain the abundance of aquatic macrophyte isotope signatures found within primary consumer tissues in our study. Across all consumer groups, all sources provided as model inputs were represented in all consumer diets, in varying proportions. This suggests that, while a preference is observed among primary consumers at each site, invertebrates are, in general, energetically supported by multiple producer groups in GRE wetlands.

## Conclusions

The goals of this study were to identify dominant energy sources supporting lower trophic levels in the GRE and to evaluate whether THg as a food web tracer could inform the source proportion estimates of food web models. Our findings suggest that wetlands in the GRE were not supported by a single dominant resource, but a dynamic mixture of detritus, macrophytes, and algae, all of which are linked to the high primary productivity in the ecosystem. Our study highlights the importance of maintaining wetland biodiversity, particularly in plant and algal communities. Although detritus was found to be a major energy resource fueling lower trophic levels, algal and macrophyte-derived carbon also contributed to primary consumers across the GRE. The digestibility and availability of algae in GRE wetlands likely contribute to its representation in consumer diets (Hart & Lovvorn; McCutchan & Lewis 2002). Similarly, the indirect contribution of submersed vegetation to odonates represents the importance of compositionally diverse aquatic plant communities in facilitating energy transfer to higher trophic levels in wetlands. Preserving this productivity and biodiversity are crucial to sustaining lower trophic levels and maintaining the structure of wetland food webs as a whole.

Because source contributions in the mixing model are based on available data, incorporating additional informative inputs such as additional tracers yields more precise posterior distribution estimates and reduces model uncertainty. Supplementing the standard dual-isotope approach clarified trophic pathways that would otherwise likely be unresolved with C and N alone due to complex cycling of these nutrients within this system. Given the variation that was observed in both δ^13^C and δ^15^N in this system, THg acted as a reliable supplement to ^15^N signals where complex N cycling is detrimental to food web resolution. In systems where primary producers can access multiple sources of N that are isotopically distinct or where anthropogenic N inputs alter baseline ^15^N exhibited among primary producers, ^15^N isotopes alone may provide inconsistent measures of trophic position. Nutrient-rich or anthropogenically impacted systems will often have these conditions and in such cases, THg may biomagnify more predictably and provide a reliable alternative indicator of trophic relationships.

## Acknowledgements

Financial support for this project included a Presidential Research Grant awarded to Alyssa M. Smith and a Catalyst Grant awarded to Dr. Matthew Cooper by Grand Valley State University. Special thanks to those who contributed to field sampling and laboratory work for this project, including Alexis Deephouse, Brenden Reid, John Gargasz, Addison Plummer, and Jenna Ulrich.

## CRediT Author Contribution Statement

**AMS:** conceptualization (equal) formal analysis, funding acquisition (supporting), methodology (lead), writing- original draft preparation. **MJC:** conceptualization (equal), funding acquisition (lead), methodology (supporting), resources (lead), supervision (lead), writing- review and editing (lead). **RRO:** methodology (supporting), resources (supporting), supervision (supporting), writing- review and editing (supporting).

## References

Albert, D. A., Wilcox, D. A., Ingram, J. W., & Thompson, T. A. (2005). Hydrogeomorphic classification for Great Lakes coastal wetlands. Journal of Great Lakes Research, 31(Suppl 1), 129–146. 10.1016/S0380-1330(05)70294-X

Anderson, O., Harrison, A., Heumann, B., Godwin, C., Uzarski, D. (2023) The influence of extreme water levels on coastal wetland extent across the Laurentian Great Lakes, Science of The Total Environment, Volume 885, 2023, 163755, ISSN 0048-9697, 10.1016/j.scitotenv.2023.163755.

Austin, J. C., Anderson, S., Courant, P. N., & Litan, R. E. (2007). America’s North Coast: A benefit–cost analysis of a program to protect and restore the Great Lakes. Brookings Institution.

Crosbie, B. & Chow-Fraser, P. (1999). Percentage land use in the watershed determines the water and sediment quality of 22 marshes in the Great Lakes basin. Canadian Journal of Fisheries and Aquatic Sciences, 56(10), 1781–1791. 10.1139/f99-109

Baattrup-Pedersen, A., Friis, K.B., Friberg, N. et al. Inter-linkages between in-stream plant diversity and macroinvertebrate communities. Hydrobiologia 852, 235–247 (2025). 10.1007/s10750-024-05700-5

Blake, W. H., Boeckx, P., Stock, B. C., Smith, H. G., Bodé, S., Upadhayay, H. R., Gaspar, L., Goddard, R., Lennard, A. T., Lizaga, I., Lobb, D. A., Owens, P. N., Petticrew, E. L., Kuzyk, Z. Z. A., Gari, B. D., Munishi, L., Mtei, K., Nebiyu, A., Mabit, L., Navas, A., … Semmens, B. X. (2018, August 30). A deconvolutional Bayesian mixing model approach for river basin sediment source apportionment. Scientific Reports, 8(1), Article 13073. 10.1038/s41598-018-30905-9

Blake, W. J., Yarnes, C. T., Cook, B. A., & James, A. C. (2011). On the use of stable isotopes in trophic ecology. Annual Review of Ecology, Evolution, and Systematics, 42, 411–440. 10.1146/annurev-ecolsys-102209-144726

Bond, A. L., & Diamond, A. W. (2011). Recent Bayesian stable-isotope mixing models are highly sensitive to variation in discrimination factors. Ecological Applications, 21(4), 1017–1023. 10.1890/09-2409.1

Brazner, J., & Sierzen, M. E., Keough, J. R., & Tanner, D. (2001). Assessing the ecological importance of coastal wetlands in a large lake context. Verhandlungen des Internationalen Vereins Limnologie, 26, 1950–1961. 10.1080/03680770.1998.11901583

Brazner, J. C., Danz, N., Niemi, G., Regal, R., Trebitz, A., Howe, R., Hanowski, J., Johnson, L., Ciborowski, J., Johnston, C., Reavie, E., Brady, V., & Sgro, G. (2007). Evaluating geographic, geomorphic, and human influences on Great Lakes wetland indicators: Multi-assemblage variance partitioning. Ecological Indicators, 7, 610–635.

Brazner, J., & Trebitz, A. (2016). Coastal wetlands of Lake Superior’s south shore (USA). In M. L. Spaans (Ed.), Coastal Wetlands: Planning and Management (pp. 235– ?). 10.1007/978-94-007-6173-5_235-1

Cabana, G., & Rasmussen, J. B. (1994). Modelling food chain structure and contaminant bioaccumulation using stable nitrogen isotopes. Nature, 372, 255–257. 10.1038/372255a0

Cabana, G., Tremblay, A., Kalff, J., & Rasmussen, J. B. (1994). Pelagic food-chain structure in Ontario lakes: A determinant of mercury levels in lake trout (*Salvelinus namaycush*). Canadian Journal of Fisheries and Aquatic Sciences, 51, 381–389. 10.1139/f94-039

Campeau, S., Murkin, H. R., & Titman, R. D. (1994). Relative importance of algae and emergent plant litter to freshwater marsh invertebrates. Canadian Journal of Fisheries and Aquatic Sciences, 51(3), 681–692. 10.1139/f94-068

Cooper, M., Uzarski, D., Burton, T., & Rediske, R. (2006). Macroinvertebrate community composition relative to chemical/physical variables, land use and cover, and vegetation types within a Lake Michigan drowned river mouth wetland. Aquatic Ecosystem Health & Management, 9, 463–479. 10.1080/14634980600892655

Cooper, M. J., Lamberti, G. A., & Uzarski, D. G. (2014). Spatial and temporal trends in invertebrate communities of Great Lakes coastal wetlands, with emphasis on Saginaw Bay of Lake Huron. Journal of Great Lakes Research, 40, 168–182.

Cooper, M.J., Uzarski, D.G. (2016). Invertebrates in Great Lakes Marshes. In: Batzer, D., Boix, D. (eds) Invertebrates in Freshwater Wetlands. Springer, Cham. 10.1007/978-3-319-24978-0_9

Craig, E. H., Arts, M. T., & Weseloh, D. V. C. (2006). [Title unavailable]. Environmental Science & Technology, 40(18), 5618–5623. 10.1021/es0520619

Dodds, W. K. (2002). Hydrology and physiography of groundwater and wetland habitats. In W. K. Dodds (Ed.), Aquatic Ecology: Freshwater Ecology (pp. 46–67). Academic Press. 10.1016/B978-012219135-0/50005-2

Diller, S., Harrison, A., Kowalski, K., Brady, V., Ciborowski, J., Cooper, M., Dumke, J., Gathman, J., Ruetz, C., Uzarski, D., Wilcox, D., & Schaeffer, J. (2022). Influences of seasonality and habitat quality on Great Lakes coastal wetland fish community composition and diets. Wetlands Ecology and Management, 30. 10.1007/s11273-022-09862-8

Eglite, E., Mohm, C., & Dierking, J. (2023). Stable isotope analysis in food web research: Systematic review and a vision for the future for the Baltic Sea macro-region. Ambio, 52(2), 319–338. 10.1007/s13280-022-01785-1

Fry, B., Bern, A. L., Ross, M. S., & Meeder, J. F. (2000). δ15N studies of nitrogen use by the red mangrove, Rhizophora mangle L. in south Florida. Estuarine, Coastal and Shelf Science, 50(2), 291–296.

Fry, B. (2002). Conservative mixing of stable isotopes across estuarine salinity gradients: A conceptual framework for monitoring watershed influences on downstream fisheries production. Estuaries, 25, 264–271. 10.1007/BF02691313

Fry, B. (2006). Stable isotope ecology. [Publisher not provided].

Garcia-Cuartas, N., Niño-Torres, C. A., Castelblanco-Martínez, D. N., Delgado-Huertas, A., Cetz-Navarro, N. P., Ortiz-Pulido, R., & Cuevas, J. (2021). Isotopic composition of aquatic and semiaquatic plants from the Mexican Caribbean: A baseline for regional ecological studies. Estuarine, Coastal and Shelf Science, 260, Article 107489. 10.1016/j.ecss.2021.107489

García-Oliva, O., & Wirtz, K. (2025). The complex structure of aquatic food webs emerges from a few assembly rules. Nature Ecology & Evolution, 9, 576–588. 10.1038/s41559-025-02647-1

Gelman, A., Carlin, J. B., Stern, H. S., & Rubin, D. B. (2014). Bayesian data analysis (3rd ed.). Taylor & Francis.

Gronewold, A. D., Fortin, V., Lofgren, B., …et al,. (2013). Coasts, water levels, and climate change: A Great Lakes perspective. Climatic Change, 120, 697–711. 10.1007/s10584-013-0840-2

Guo, F., Kainz, M.J., Sheldon, F. and Bunn, S.E. (2016), The importance of high-quality algal food sources in stream food webs – current status and future perspectives. Freshw Biol, 61: 815–831. 10.1111/fwb.12755

Handley, L. L., & Raven, J. A. (1992). The use of natural abundance of nitrogen isotopes in plant physiology and ecology. Plant, Cell & Environment, 15, 965–985. 10.1111/j.1365-3040.1992.tb01650.x

Hart, C., & Kilham, S. S. (2004). Elevated δ15N in stream biota in areas with septic tank systems in an urban watershed. Ecological Applications, 14, 637–641. 10.1890/03-5148

Hart, E. A., & Lovvorn, J. R. (2003). Algal vs. macrophyte inputs to food webs of inland saline wetlands. Ecology, 84, 3317–3326. 10.1890/02-0642

Herdendorf, C. E. (1987). The ecology of Lake Erie coastal marshes: A community profile (Biol. Rep. No. 85(7.9)). U.S. Fish & Wildlife Service.

Herdendorf, C. E. (1990). Great Lakes estuaries. Estuaries, 13(4), 493–503. 10.2307/1351795

Jeffres CA, Holmes EJ, Sommer TR, Katz JVE (2020) Detrital food web contributes to aquatic ecosystem productivity and rapid salmon growth in a managed floodplain. PLOS ONE 15(9): e0216019. 10.1371/journal.pone.0216019

Kelly, J. R., & Scheibling, R. E. (2012). Fatty acids as dietary tracers in benthic food webs. Marine Ecology Progress Series, 446, 1–22. 10.3354/meps09559

Keough, J. R., Thompson, T. A., Guntenspergen, G. R., & Wilcox, D. A. (1999). Hydrogeomorphic factors and ecosystem responses in coastal wetlands of the Great Lakes. Wetlands, 19, 821–834. 10.1007/BF03161786

King, R. S., & Brazner, J. C. (1999). Coastal wetland insect communities along a trophic gradient in Green Bay, Lake Michigan. Wetlands, 19, 426–437. 10.1007/BF03161774

Krumins, J. A., van Oevelen, D., Bezemer, T. M., De Deyn, G. B., Hol, W. G., van Donk, E., Van der Putten, W. H. (2013). Soil and freshwater and marine sediment food webs: Their structure and function. BioScience, 63(1), 35–42.

Larson, J. H., Trebitz, A. S., Steinman, A. D., Wiley, M. J., Mazur, M. C., Pebbles, V., Braun, H. A., & Seelbach, P. W. (2013). Great Lakes rivermouth ecosystems: Scientific synthesis and management implications. Journal of Great Lakes Research, 39(3), 513–524. 10.1016/j.jglr.2013.06.002

Lavoie, R. A., Jardine, T. D., Chumchal, M. M., Kidd, K. A., & Campbell, L. M. (2013). Biomagnification of mercury in aquatic food webs: A worldwide meta-analysis. Environmental Science & Technology, 47(23), 13385–13394. 10.1021/es403103t

Layer, K., Hildrew, A.G. & Woodward, G. Grazing and detritivory in 20 stream food webs across a broad pH gradient. Oecologia 171, 459–471 (2013). 10.1007/s00442-012-2421-x

Lichtfouse E, Lichtfouse M, Jaffrézic A. Delta13C values of grasses as a novel indicator of pollution by fossil-fuel-derived greenhouse gas CO2 in urban areas. Environ Sci Technol. 2003 Jan 1;37(1):87–9. doi: 10.1021/es025979y. PMID: 12542295.

Lindeman, R. L. (1942). The trophic-dynamic aspect of ecology. Ecology, 23, 399–417. 10.2307/1930126

Luoma, S. N., & Rainbow, P. S. (2008). Metal contamination in aquatic environments: science and lateral management (Vol. 126). Cambridge: Cambridge university press.

Ma, M., Du, H., & Wang, D. (2019). Mercury methylation by anaerobic microorganisms: A review. Critical Reviews in Environmental Science and Technology, 49(20), 1893–1913.

Mader, M. M., Ruetz, C., Woznicki, S. A., Steinman, A. D., Land cover and water quality of drowned river mouths: Evidence of an environmental gradient along the eastern Lake Michigan shoreline, Journal of Great Lakes Research, Volume 49, Issue 6, 2023, 102237, ISSN 0380-1330, 10.1016/j.jglr.2023.09.008.

Marty, J., & Planas, D. (2008). Comparison of methods to determine algal δ¹³C in freshwater. Limnology and Oceanography: Methods, 6, 51–63.

McClelland, J. W., & Valiela, I. (1998). Linking nitrogen in estuarine producers to land-derived sources. Limnology and Oceanography, 43(4), 577–585. 10.4319/lo.1998.43.4.0577

McClelland, J. W., Valiela, I., & Michener, R. H. (1997). Nitrogen-stable isotope signatures in estuarine food webs: A record of increasing urbanization in coastal watersheds. Limnology and Oceanography, 42, 930–940. 10.4319/lo.1997.42.5.0930

McCutchan, J. H., Jr., & Lewis, W. M., Jr. (2002). Relative importance of carbon sources for macroinvertebrates in a Rocky Mountain stream. Limnology and Oceanography, 47(3), 742–748. 10.4319/lo.2002.47.3.0742

Michigan PFAS Action Response Team. (n.d.). J.B. Sims Generating Station. Michigan Department of Environment, Great Lakes, and Energy. https://www.michigan.gov/pfasresponse/investigations/sites-aoi/ottawa-county/jb-sims-generating-station

Middelburg, J. J. (2014). Stable isotopes dissect aquatic food webs from the top to the bottom. Biogeosciences, 11, 2357–2371. 10.5194/bg-11-2357-2014

Midwood, J. D., & Chow-Fraser, P. (2012). Changes in aquatic vegetation and fish communities following 5 years of sustained low water levels in coastal marshes of eastern Georgian Bay, Lake Huron. Global Change Biology, 18(1), 93–105.

Moore, J. C., Berlow, E. L., Coleman, D. C., De Ruiter, P. C., Dong, Q., Hastings, A., … & Wall, D. H. (2004). Detritus, trophic dynamics and biodiversity. Ecology letters, 7(7), 584–600.

Nieoczym, M., Stryjecki, R., Buczyński, P. et al. Differential abundance, composition and mesohabitat use by aquatic macroinvertebrate taxa in ponds with and without fish. Aquat Sci 85, 25 (2023). 10.1007/s00027-022-00922-y

Nogues, Q., Baulaz, Y., Clavel, J., Araignous, E., Bourdaud, P., Ben Rais Lasram, F., Dauvin, J.-C., Girardin, V., Halouani, G., Le Loc’h, F., Loew-Turbout, F., Raoux, A., & Niquil, N. (2023). The usefulness of food web models in the ecosystem services framework: Quantifying, mapping, and linking services supply. Ecosystem Services, 63, 101550. 10.1016/j.ecoser.2023.101550

Odum, E. P. (1971). Fundamentals of ecology (3rd ed.). W. B. Saunders.

Olsen, Y. S., Fox, S. E., Kinney, E. L., Teichberg, M., & Valiela, I. (2010). Differences in urbanization and degree of marine influence are reflected in δ¹³C and δ¹⁵N of producers and consumers in seagrass habitats of Puerto Rico. Marine Environmental Research, 69(3), 198–206. 10.1016/j.marenvres.2009.10.005

Peterson, B. J., & Fry, B. (1987). Stable isotopes in ecosystem studies. Annual Review of Ecology and Systematics, 18, 293–320. http://www.jstor.org/stable/2097134

Phillips, D. L., Inger, R., Bearhop, S., Jackson, A. L., Moore, J. W., Parnell, A. C., Semmens, B. X., & Ward, E. J. (2014). Best practices for use of stable isotope mixing models in food-web studies. Canadian Journal of Zoology, 92(10), 823–835.

Post, D. M. (2002). Using stable isotopes to estimate trophic position: Models, methods, and assumptions. Ecology, 83, 703–718. 10.1890/0012-9658(2002)083[0703:USITET]2.0.CO;2

Purdy K.J., Hurd P.J., Moya-Larano J., Trimmer M., Oakley B.B. & Woodward G. (2010) Systems biology for ecology:from molecules to ecosystems. Advances in EcologicalResearch, 43, 87–149

R Core Team. (2024). R: A language and environment for statistical computing. R Foundation for Statistical Computing.

Ratkowsky, D. A., Dix, T. G., & Wilson, K. C. (1975). Mercury in fish in the Derwent Estuary, Tasmania, and its relation to the position of the fish in the food chain. Australian Journal of Marine and Freshwater Research, 26, 223–231. 10.1071/MF9750223

Rojas, T.V., O’Reilly, K.E., Houghton, C.J., Shrovnal, J.S., Berg, M.B., Uzarski, D.G., Lamberti, G.A. and Forsythe, P.S. (2025), Coastal Wetlands Drive Isotopic Niche Plasticity of Top Predator Fish Communities in Green Bay, Lake Michigan (USA). Ecol Evol, 15: e71463. 10.1002/ece3.71463

Schlesinger, W. H., & Bernhardt, E. S. (2020). Wetland ecosystems. In W. H. Schlesinger & E. S. Bernhardt (Eds.), Biogeochemistry (4th ed., pp. 249–291). Academic Press. 10.1016/B978-0-12-814608-8.00007-4

Sierszen, M. E., Brazner, J. C., Cotter, A. M., Morrice, J. A., Peterson, G. S., & Trebitz, A. S. (2012). Watershed and lake influences on the energetic base of coastal wetland food webs across the Great Lakes Basin. Journal of Great Lakes Research, 38(3), 418–428. 10.1016/j.jglr.2012.04.005

Sierszen, M. E., Schoen, L. S., Kosiara, J. M., Hoffman, J. C., Cooper, M. J., & Uzarski, D. G. (2019). Relative contributions of nearshore and wetland habitats to coastal food webs in the Great Lakes. Journal of Great Lakes Research, 45(1), 129–137. 10.1016/j.jglr.2018.11.006

Sinclair, C. A., Garcia, T. S., & Eagles-Smith, C. A. (2024). A meta-analysis of mercury biomagnification in freshwater predatory invertebrates: Community diversity and dietary exposure drive variability. Environmental Science & Technology, 58(43), 19429–19439. 10.1021/acs.est.4c05920

Soto, D., Gacia, E., & Catalan, J. (2013). Freshwater food web studies: A plea for multiple tracer approach. Limnetica, 32, 97–106.

Stephens, R. B., Shipley, O. N., & Moll, R. J. (2023). Meta-analysis and critical review of trophic discrimination factors (Δ¹³C and Δ¹⁵N): Importance of tissue, trophic level and diet source. Functional Ecology, 37, 2535–2549. 10.1111/1365-2435.14403

Steffy, L. Y., & Kilham, S. S. (2004). Elevated δ¹⁵N in stream biota in areas with septic tank systems in an urban watershed. Ecological Applications, 14, 637–641. 10.1890/03-5148

Stock, B. C., & Semmens, B. X. (2016). *MixSIAR GUI user manual* (Version 3.1). 10.5281/zenodo.1209993

Sutherland, W. J., et al. (2013). Identification of 100 fundamental ecological questions. Journal of Ecology, 101, 58–67. 10.1111/1365-2745.12025

Teal, J. M. (1962). Energy flow in the salt marsh ecosystem of Georgia. Ecology, 43, 614–624. 10.2307/1933451

Thompson, R. M., Brose, U., Dunne, J. A., Hall, R. O., Hladyz, S., Kitching, R. L., Martinez, N. D., Rantala, H., Romanuk, T. N., Stouffer, D. B., & Tylianakis, J. M. (2012). Food webs: Reconciling the structure and function of biodiversity. Trends in Ecology & Evolution, 27(12), 689–697. 10.1016/j.tree.2012.08.005

Thompson, R. M., Dunne, J. A., & Woodward, G. (2012). Freshwater food webs: Towards a more fundamental understanding of biodiversity and community dynamics. Freshwater Biology, 57, 1329–1341. 10.1111/j.1365-2427.2012.02808.x

James H. Thorp, D. Christopher Rogers, Alan P. Covich, Chapter 27 - Introduction to “Crustacea”, Editor(s): James H. Thorp, D. Christopher Rogers, Thorp and Covich’s Freshwater Invertebrates (Fourth Edition) Academic Press, 2015, Pages 671–686, ISBN 9780123850263, 10.1016/B978-0-12-385026-3.00027-9.

Trebitz, A. S., Cotter, A. M., & Morrice, J. A. (2002). Relative role of lake and tributary in hydrology of Lake Superior coastal wetlands. Journal of Great Lakes Research, 28, 212–227.

Trebitz, A.S., Morrice, J.A., Taylor, D.L. et al. Hydromorphic determinants of aquatic habitat variability in Lake Superior coastal wetlands. Wetlands 25, 505–519 (2005). 10.1672/0277-5212(2005)025[0505:HDOAHV]2.0.CO;2

Trebitz, A. S. (2006). Characterizing seiche and tide-driven daily water level fluctuations affecting coastal ecosystems of the Great Lakes. Journal of Great Lakes Research, 32(1), 102–116.

Uzarski, D. G., Burton, T. M., Cooper, M. J., Ingram, J., & Timmermans, S. (2005). Fish habitat use within and across wetland classes in coastal wetlands of the five Great Lakes: Development of a fish-based index of biotic integrity. Journal of Great Lakes Research, 31(1), 171–187.

Uzarski, D. G., Burton, T. M., Kolar, R. E., & Cooper, M. J. (2009). The ecological impacts of fragmentation and vegetation removal in Lake Huron’s coastal wetlands. Aquatic Ecosystem Health & Management, 12(1), 45–62. 10.1080/14634980802690881

Valiela, I., Foreman, K., LaMontagne, M., et al. (1992). Couplings of watersheds and coastal waters: Sources and consequences of nutrient enrichment in Waquoit Bay, Massachusetts. Estuaries, 15, 443–457. 10.2307/1352389

van der Merwe, J., & Hellgren, E. C. (2016). Spatial variation in trophic ecology of small mammals in wetlands: Support for hydrological drivers. Ecosphere, 7(11), Article e01567. 10.1002/ecs2.1567

Vander Zanden, Jake & Rasmussen, Joseph. (2001). Variation in δ15N and δ13C trophic fractionation. Limnology and Oceanography. 46. 2061–2066. 10.4319/lo.2001.46.8.2061.

Wetzel, R. G. (1992). Wetlands as metabolic gates. Journal of Great Lakes Research, 18(4), 529–532. 10.1016/S0380-1330(92)71320-3

Wickham, H. (2016). ggplot2: Elegant graphics for data analysis. Springer-Verlag New York.

Wilcox, D. A. (2004). Implications of hydrologic variability on the succession of plants in Great Lakes wetlands. Aquatic Ecosystem Health & Management, 7(2), 223–231. 10.1080/14634980490461579

Wissinger, S. A. (1999). Ecology of wetland invertebrates: Synthesis and applications for conservation and management. In D. P. Batzer, R. D. Rader, & S. A. Wissinger (Eds.), Invertebrates in Freshwater Wetlands of North America: Ecology and Management (pp. 1043–1086). Wiley.

