## Supplemental Information for "Advancing Great Lakes Coastal Wetland Food Web Models using an Integrative Tracer Approach"

Table S1: Isotopic ratios for ^13^C and ^15^N, and total mercury (THg) concentrations in µg/kg (mean±SE) across sites and sample types.

|  | Site | Algae | SAV | Macrophytes | Detritus | *Amphipoda* | *Odonata* | Phytoplankton | Seston | Zooplankton |
| --- | --- | --- | --- | --- | --- | --- | --- | --- | --- | --- |
| THg (µg/kg) | PET | 7.896± 1.720 | 12.94± 7.65 | 4.945± 2.356 | 93.497± 28.546 | 30± 0 | 73.486± 13.913 |  |  |  |
|  | BR1 | 4.931± 1.0 | 8.574± 2.138 | 3.617± 0.214 | 146.104± 24.431 | 40± 0 | 24.810± 17.185 |  |  |  |
|  | STR | 7.126± 1.495 | 15.935± 4.148 | 2.768± 0.365 | 155.180± 9.579 | 27.50± 3.535 | 88.143± 45.845 |  |  |  |
|  | PT1 | 15.736± 3.63 | 8.298± 1.134 | 4.038± 0.786 | 125.593± 63.179 | 46.25± 0 | 75.190± 21.215 |  |  |  |
|  | SGA | 4.409± 0.363 | 6.456± 1.350 | 2.015± 0.551 | 107.015± 4.574 | 11.111± 0 | 38.597± 19.622 |  |  |  |
|  | HI2 | 7.764± 0.20 | 5.870± 1.120 | 1.944± 0.304 | 119.045± 26.732 | 22.903± 0 | 57.779± 29.76 |  |  |  |
| δ^13^C | PET | -18.406± 3.085 | -29.613± 2.344 | -26.64± 0.648 | -28.103± 0.824 | -25.397± 2.17 | -28.700± 0.481 | -27.366± 1.151 | -27.993± 0.521 | -26.223± 0.426 |
|  | BR1 | -29.170± 1.394 | -28.066± 3.836 | -25.49± 0.626 | -28.270± 0.305 | -28.778± 0.363 | -30.798± 0.142 | -29.202± 0.140 | -30.156± 0.171 | -29.535± 0.780 |
|  | STR | -27.593± 1.697 | -36.413± 1.668 | -26.216± 0.318 | -28.493± 0.136 | -28.258± 0.244 | -31.218± 1.886 | -28.138± 1.885 | -29.445± 0.532 | -30.207± 1.596 |
|  | PT1 | -29.603± 3.188 | -32.296± 3.748 | -26.836± 0.494 | -30.218± 0.408 | -32.217± 0.563 | -33.578± 1.108 | -26.723± 1.084 | -29.928± 1.014 | -29.933± 1.667 |
|  | SGA | -24.073± 3.366 | -26.146± 2.202 | -24.726± 0.357 | -22.685± 2.707 | -24.159± 0.566 | -26.332± 0.926 | -22.533± 0.617 | -23.566± 0.642 | -26.078± 0.767 |
|  | HI2 | -21.01± 2.340 | -26.813± 1.719 | -25.91± 0.892 | -23.041± 0.746 | -25.877± 0.206 | -27.896± 0.809 | -21.697± 0.373 | -21.933± 0.833 | -25.704± 1.044 |
| δ^15^N | PET | 10.453± 0.328 | 9.103± 0.610 | 4.456± 2.706 | 3.217± 0.068 | 8.592± 0.522 | 9.952± 1.245 | 5.068± 0.629 | 6.164± 0.989 | 7.941± 0.388 |
|  | BR1 | 6.080± 1.207 | 6.373± 1.240 | 4.853± 0.782 | 3.019± 0.473 | 5.499± 0.805 | 7.335± 1.343 | 4.924± 0.060 | 3.983± 0.099 | 4.148± 0.755 |
|  | STR | 2.736± 1.102 | 5.886± 0.335 | 4.313± 1.053 | 2.606± 0.193 | 4.587± 0.042 | 6.481± 1.245 | 5.360± 1.146 | 3.547± 0.275 | 2.822± 1.159 |
|  | PT1 | 5.033± 1.60 | 5.130± 0.771 | 4.673± 0.720 | 3.275± 0.175 | 5.695± 0.246 | 7.305± 0.091 | 5.511± 0.238 | 3.941± 0.458 | 4.570± 0.629 |
|  | SGA | 11.713± 0.713 | 11.406± 1.072 | 4.25± 1.480 | 6.393± 0.171 | 10.740± 0.244 | 12.410± 0.438 | 7.711± 0.838 | 6.068± 2.20 | 10.290± 0.670 |
|  | HI2 | 10.853± 0.406 | 12.840± 0.694 | 6.926± 1.60 | 6.271± 0.090 | 11.194± 0.458 | 12.654± 0.894 | 7.80± 0.311 | 8.520± 0.163 | 10.712± 0.274 |
